# Exceptional ancient DNA preservation and fibre remains of a Sasanian saltmine sheep mummy in Chehrābād, Iran

**DOI:** 10.1101/2021.04.15.439892

**Authors:** Conor Rossi, Gabriela Ruß-Popa, Valeria Mattiangeli, Fionnuala McDaid, Andrew J. Hare, Hossein Davoudi, Haeedeh Laleh, Zahra Lorzadeh, Roya Khazaeli, Homa Fathi, Matthew D. Teasdale, Abolfazl A’ali, Thomas Stöllner, Marjan Mashkour, Kevin G. Daly

## Abstract

Mummified remains have long attracted interest as a potential source of ancient DNA. However, mummification is a rare process that requires an anhydrous environment to rapidly dehydrate and preserve tissue before complete decomposition occurs. We present the whole genome sequences of a ∼1600 year old naturally mummified sheep recovered from Chehrābād, a salt mine in northwestern Iran. Comparative analyses of published ancient sequences revealed remarkable DNA integrity of this mummy. Hallmarks of postmortem damage, fragmentation and hydrolytic deamination, are substantially reduced, likely due to the high-salinity of this taphonomic environment. Metagenomic analyses reflect the profound influence of high salt content on decomposition; its microbial profile is predominated by halophilic archaea and bacteria, possibly contributing to the preservation of this sample. Applying population genomic analyses we find consistent clustering of this sheep with Southwest Asian modern breeds, suggesting ancestry continuity. Genotyping of a locus influencing the woolly phenotype showed the existence of an ancestral “hairy” allele in this sheep, consistent with hair fibre imaging, further elucidating Sasanian-period animal husbandry.

## Introduction

In 1993, a remarkably preserved human body dating to the ∼1700 years Before Present (BP) was discovered in the Douzlākh salt mine near Chehrābād village in the Zanjan Province of northwest Iran [1–3]. A total of 8 “Salt Men” have been identified at the mine [4,5], several retaining keratinous tissues such as skin, hair, and both endo- and exoparasites, despite dating to the Achaemenid (550-330 BCE, 2500-2280 BP) and Sasanian (224-651 CE, ∼1700-1300 BP) periods. The mine, also known as Chehrābād, was active in various periods and its archaeological refilling layers represent an extraction history that ranged from the 6^th^ century BCE to 20^th^ century CE. In addition to the “Salt Men”, textiles, leather objects, and animal remains have been discovered [6,7], likely preserved by high salinity and low moisture content of the mine. Isotopic, genetic, and lipid analyses have been reported for this material [1], and studies have been carried out to characterize genomic DNA survival [8]. These human and animal remains are examples of natural mummification -the spontaneous desiccation of soft tissue by a dry environment that rapidly dehydrates soft tissue before decay begins [9].

Mummification has been suggested as a mechanism that may sufficiently preserve keratinised tissue for ancient DNA (aDNA) sequencing [9]. The effects of age-related damage in aDNA are well documented and include base misincorporation at strand overhangs, fragmentation and low endogenous content [10]. Both deamination and depurination, associated with postmortem transition error and DNA fragmentation, respectively, require water as a substrate [11]. Ancient DNA from Chehrābād, a highly saline, anhydrous environment, presents an opportunity to investigate potential differences in nucleotide degradation resulting from this unusual taphonomic context.

In this study we sequenced DNA from the ∼1600 year old (Sasanian period) mummified sheep leg 4305, recently discovered in a large mining gallery in the northwestern edge of the Douzlākh saltmine of Chehrābād by Iranian-German researchers during archaeological excavations (Fig. 1A) [2]. The specimen was likely deposited during refilling activities in the 4-5th centuries CE after the gallery’s reopening in the Early Sasanian period (2nd-3rd centuries CE) and following its initial collapse between 405-380 BCE. The leg was possibly discarded during food preparation activities, as both sheep and goat were likely used as provisioning for Sasanian-period miners; equines may have been used as beasts of burden [12]. We find unusual survival patterns of endogenous DNA given its distance from the equator, implying exceptional preservation of nucleic acid integrity was afforded by the unique salt-rich environment. This enables population genomic profiling of this sheep including genotyping the antisense *EIF2S2* retrogene insertion within the 3’ UTR of the *IRF2BP2* gene which influences the woolly phenotype [13], in tandem with fibre analysis using scanning electron microscopy (SEM), and characterising the mummy skin metagenome.

**Figure 1.**
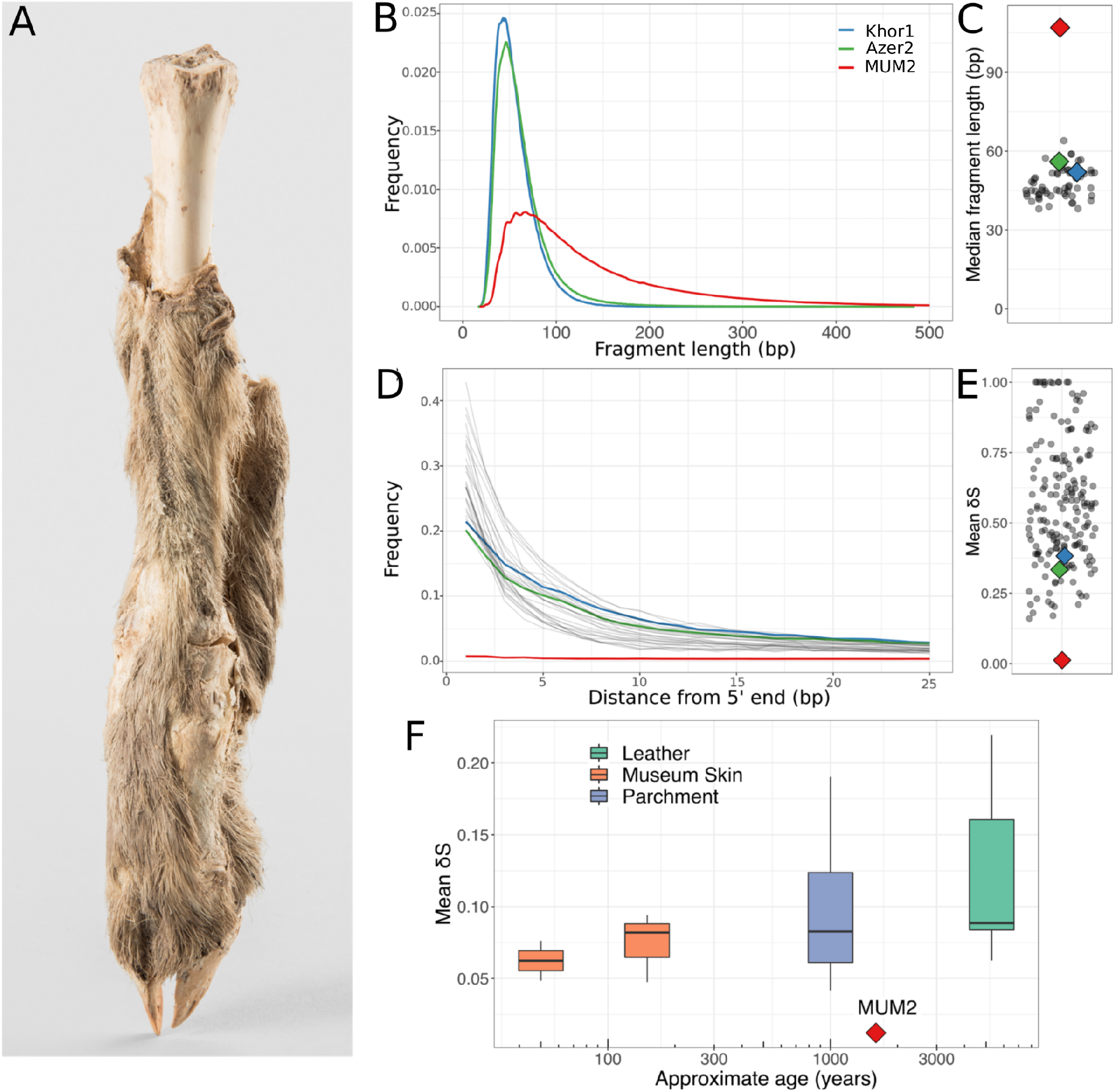
(A) Mummified sheep leg (4305) after cleaning. Photography: N. Tehrani **(B) Read length distributions of MUM2, Khor1, and Azer2, calculated from PE data**. MUM2 shows a reduced rate of fragmentation. The median read length of MUM2 (107 bp) exceeds the median read length of Khor1 and Azer2 (52 bp & 56 bp, respectively). **(C) Median read lengths of 61 published ancient Ovicaprid samples** [50]. The median read length of MUM2 (107 bp; 90 bp among collapsed reads only) exceeds the longest among published ovicaprid genomes (64 bp). **(D) Deamination patterns of MUM2, Khor1, Azer2, and other ancient ovicaprids for non UDG-treated libraries**. Low levels of base misincorporation at the 5’ ends of reads were observed for MUM2 compared to Khor1 and Azer2. **(E) Mean** δ**S of published 182 ancient bone samples** [50] [41]. The mean δS of MUM2 (0.012) is singular in its low levels of deamination. **(F) Comparison of mean** δ**S of published ancient skins** [51] [49] **[48]**. **Lower damage rates are recorded compared to all samples, including some ∼50 years old**.

## Materials & Methods

A sample of the mummified sheep skin (Table 1, MUM2) from sheep leg 4305 was directly radiocarbon dated at the ^**14**^CHRONO Centre (Queen’s University Belfast). OxCal 4.3.2 [14] was used to calibrate its age (95.4% confidence interval) using [15].

**Table 1.** Summary information of samples sequenced in this study.

| Name | Tissue | Origin | Period | Age | Endogenous DNA % | Coverage (X) |
| --- | --- | --- | --- | --- | --- | --- |
| MUM2 | Mummified skin | Chehrābād, Iran | Sasanian Empire period | *399-539 cal CE | 31.01 | 3.94 |
| Khor1 | Petrous bone | Nishapur, Iran | Sasanian - Islamic periods | 600 - 1200 CE | 58.44 | 0.04 |
| Azer2 | Petrous bone | Tepe Hasanlu, Iran | Iron Age III | 800-600 BCE | 31.32 | 0.07 |

Sample preparation, extraction and library preparation were performed in a dedicated aDNA laboratory in the Smurfit Institute of Genetics, Trinity College Dublin according to standard protocols (supplementary material). Sequencing of MUM2 and two Iranian sheep bone samples (Khor1 and Azer2) of approximately similar age (Table 1) for comparison was performed on Illumina MiSeq (50bp SE) and HiSeq 2500 platforms (100bp SE and 100bp PE).

Sequencing reads were aligned to OviAri3.1 and filtered to produce bam files following standard aDNA sequencing pipelines (supplementary material). Damage patterns were assessed using mapDamage2.0 [16].

Filtered reads not aligned to either sheep or human genomes were taxonomically assigned using the metagenomic classifier Kraken2 [17]. Microbial sources were estimated using SourceTracker2 [18] with a custom metagenomic database [19] [20] [21] [22] [23] (supplementary material). Bacterial species abundances were generated using MIDAS [24].

Mitochondrial sequences were produced using ANGSD [25] and a Maximum-Likelihood phylogenetic tree was generated using SeaView and phyML [26–28] with the HKY85 substitution model, selected using jmodeltest2 [29,30] and 100 bootstrap repeats.

A SNP dataset of modern breeds [31] was used to investigate genomic affinities (supplementary material). LASER (v2.03) PCA [32], outgroup *f*_*3*_ statistics [33], TreeMix [34] and ADMIXTURE [35] analyses were completed (supplementary material).

We investigated the woolly locus located on chromosome 25 [13]. Two modified OviAri3.1 assemblies were produced, one representing the ancestral “hairy” phenotype, the other representing the “woolly” phenotype (supplementary material). Final bam files were visualised using IGV [36]. Hair fibres were examined using scanning electron and light microscopes at USTEM, TU Wien and Austrian Archaeological Institute respectively.

## Results and Discussion

The Chehrābād mummy sample (MUM2) was directly dated to the 5-6th century CE (2 sigma 1621-1481 cal BP, uncalibrated 1600 ± 30 BP, Supplementary Figure 3). This aligns with the Sasanian Empire period of Iran, a time when the mine was in active use [1]. Initial DNA screening indicated high endogenous DNA for MUM2, and also the comparative Iranian sheep samples from relatively close time periods (Table 1).

Sequencing of the Chehrābād mummy produced a 3.94X genome after quality filtering (Supplementary Table 1), in addition to the low coverage comparative genomes (0.04X and 0.07X). MUM2 differs from the two comparative sheep samples in displaying longer fragment lengths (median 107bp vs 52bp and 56bp; Figure 1B; collapsed reads-only 90bp vs 50bp and 55bp) and substantially lower rates of deamination (Figure 1D) (δS, single strand cytosine deamination probability, mean δS = 0.012 vs. 0.382 and 0.334). Contrasting previously published ancient ovicaprid data from Southwest Asia and Europe (Supplementary Table 3), MUM2 falls outside the ranges of both median fragment length and mean δS values (Figure 1C and 1E), indicating remarkably low fragmentation and deamination of the Chehrābād sheep mummy genomic material given its latitude. Similar length distributions have been reported primarily from high latitude and permafrost environments [37–40]. A low level of thermal fluctuations may also contribute to DNA preservation [41], as comparable fragment lengths have been reported in a human sample from Wezmeh Cave, Iran [42].

Recent models of postmortem DNA fragmentation suggest rate-constant hydrolytic depurination over time [43], or age-independent, driven by environment-dependent biotic and abiotic factors [41]. The depurination rates of MUM2 are similar to the more-fragmented comparative samples (Supplementary Figure 4), implying that other processes in the Chehrābād environment underlie the lower fragmentation rates. The highly alkaline, cool and anhydrous conditions may have contributed to inhibition of cellular nucleases which would otherwise degrade and fragment endogenous DNA [9]. Postmortem DNA deamination via cytosine hydrolysis [44] is thought to be strongly correlated with age [45] and thermal age [41]. The substantially-lower rates of deamination observed in MUM2 is likely due to the scarcity of environmental free water, required for hydrolytic deamination. These results are consistent with Chehrābād providing a taphonomic environment conducive to genome preservation.

DNA preservation may also be influenced by its tissue-of-origin; for example, bone hydroxyapatite rather than keratin fractions are associated with smaller fragment size [46]. As hydrophobic keratinised tissue may provide resistance to environmental water [47], we compared MUM2 to published ancient skins genomes (Figure 1F) to determine if tissue providence was solely responsible for DNA preservation. The mean δS of MUM2 falls outside the range of other ancient skin genomes, including 20th century CE goat skins [48] and leather recovered from the Tyrolean Iceman [49]. While this does not discount keratinized tissue being specifically enriched with longer DNA fragments, the Chehrābād sheep mummy appears to be singular in its DNA integrity among published skin samples.

Given the distinctive geochemical composition of Chehrābād, we examined if its salt-rich environment was reflected in the metagenomic profile of MUM2. Taxonomic assignment and abundance estimation assigned 57.13% of classified reads to the halophilic Class of Archaea *Halobacteria* (Supplementary Table 2). Similarly, SourceTracker2 predicted that 0.4725 -0.7458 of the microbial community originated from a salt-rich environment (Table 2, Supplementary Figure 5). A complementary analysis using MIDAS identified 76 unique bacterial species in the mummified sheep (Supplementary Table 4). The most abundant species is the halophilic bacterium *Actinopolyspora halophila 58532*, accounting for ∼29% of identified reads. This signal of a dominant halophilic microbial community is not replicated in comparison samples or controls (Table 2, supplementary material). Rapid colonization by saprophytic microbial communities, with key decomposers being ubiquitous across soil types, is typical for mammalian corpses post-mortem [52]. The halophilic metagenome profile observed in the Chehrābād sheep mummy skin indicates that the typical decomposers may be less abundant in this alkaline, salt-rich setting, which may have contributed to soft tissue and molecular preservation.

**Table 2:** Predicted source proportion of metagenomic reads by SourceTracker2.

|  | **MUM2**<br>**(Species)** | **Azer2**<br>**(Species)** | **Khor1**<br>**(Species)** | **MUM2**<br>**(Genus)** | **Azer2**<br>**(Genus)** | **Khor1**<br>**(Genus)** |
| --- | --- | --- | --- | --- | --- | --- |
| **Tissue decomposers** | 0.0212 | 0.1458 | 0.0386 | 0.0426 | 0.3524 | 0.3727 |
| **Salt-rich** | 0.4725 | 0.0026 | 0.0036 | 0.7458 | 0.0145 | 0.0151 |
| **Laboratory reagents** | 0.0003 | 0.0002 | 0.0002 | N/A | N/A | N/A |
| **Sheep skin** | 0.0285 | 0.085 | 0.0463 | 0.0829 | 0.3294 | 0.2244 |
| **Soil** | 0.0009 | 0.0007 | 0.0046 | 0 | 0 | 0 |
| **Unknown** | 0.4766 | 0.7657 | 0.9067 | 0.1287 | 0.3037 | 0.3878 |

### Population genomics

We investigated how the Chehrābād sheep MUM2 relates to modern populations using mitochondrial and autosomal variation. A 664X mitochondrial genome of MUM2 falls within the C haplotype cluster in a Maximum-Likelihood phylogeny of modern sheep mitochondria (Supplementary Figure 7). This clade is found at its highest frequency in southwest and east Asia [53,54], and has been reported in ancient samples from Bronze Age Turkey [55], and is consistent with past and present-day patterns of mitochondrial diversity.

PCA from autosomal variation clusters MUM2 with modern southwest Asian breeds, using both global and Asian reference panels (Supplementary Figure 8). *f*_*3*_ outgroup statistics show MUM2 shares the most genetic drift with southwest Asian breeds, particularly those from Iran (Figure 2A). ADMIXTURE and TreeMix analysis also confirmed the affinity of MUM2 with modern sheep breeds from southwest Asia, particularly the Qezel and Afec-Assaf breeds sampled from Iran [31] (supplementary material). Overall, there is genetic continuity between west Iranian sheep populations in Sassanid and modern time periods, although PCA using Ovine SNP50 genotypes of Asian breeds place MUM2 apart from sampled breeds (Supplementary Figure 9), suggesting a degree of genetic flux during the past 1500-1600 years in Iranian sheep.

**Figure 2.**
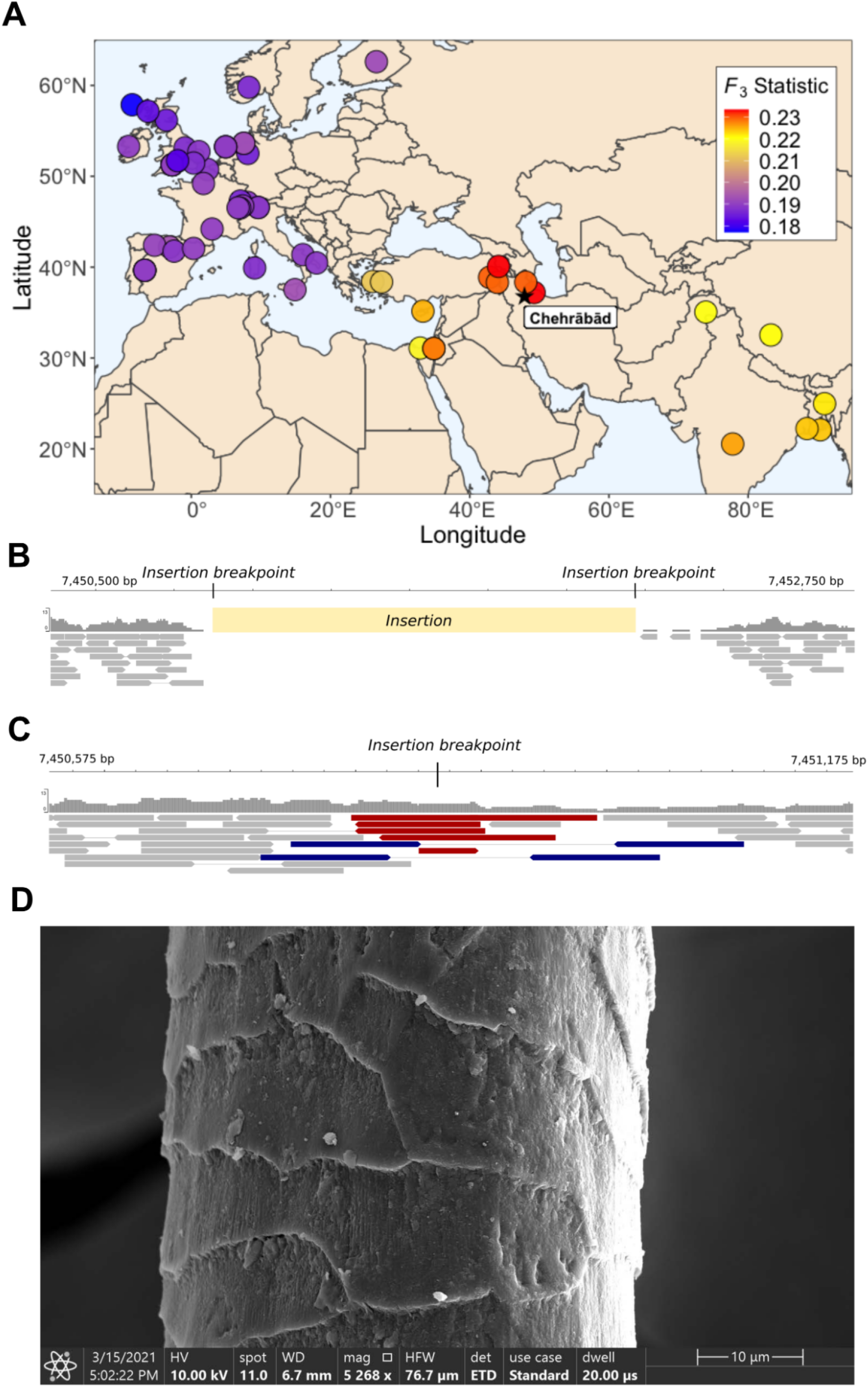
(A) Shared genetic drift between MUM2 and modern sheep populations. Higher *f*_*3*_ values, in red, indicate higher shared drift, relative to the outgroup Asiatic Mouflon. **Visualisation of read coverage of filtered bam files at woolly locus, in assemblies with (B) and without (C) the insertion**. Reads highlighted in red overlap the insertion breakpoint, blue indicates an inferred overlap of the insertion point by the straddling read pair. Highlighted reads map only to one assembly and do not align to the other. (**D) SEM image of MUM2 hair fibre, displaying the mosaic scales typical of a sheep hair shaft**. Image by A. Steiger-Thirsfeld and G. Ruß-Popa.

### Fibre Genotype and Phenotype

The derived “woolly” coat phenotype is thought to be influenced by a ∼1.5 kbp insertion of a *EIF2S2* retrogene into the *IRF2BP2* 3’ UTR, recessive to the ancestral allele associated with “hairy” coat [13]. We exploited the length of the MUM2 DNA fragments to investigate this “woolly” locus by searching for read pairs which either encompassed or overlapped the insertion breakpoint, indicative of a copy of the “hairy” allele. No reads were found to overlap the diagnostic insertion breakpoints of the “woolly” insert, which would indicate a copy of the “woolly” allele (Figure 2B). Five reads were found to uniquely map to the ‘hairy’ allele diagnostic position, with a further two read pairs inferred to overlap this breakpoint (Figure 2C). We therefore infer this animal to be either homozygous or heterozygous for the dominant “hairy” allele. In addition, SEM imaging of the mostly-unpigmented mummified hair fibres revealed mosaic scales typical of sheep [56] with fine lines on the scale surface (Figure 2D, supplementary material), a characteristic of sheep hair fibres and particularly for mouflon and medium-wool breeds [57]. This may reflect MUM2 coming from a herd maintained for meat or milk production rather than wool, consistent with suggestions that ovicaprids were used as food for workers, and that sections of the mine were used as stables [1].

The Chehrābād saltmine provides a remarkable case study of the importance of taphonomic context for tissue and genetic preservation. We exploit this to investigate both the hair fibre genotype and phenotype of this natural sheep mummy, thus intimating at its likely use as a food source for Sassian-period miners.

## Data Accessibility

Sequencing reads and mitochondrial sequences are available under European Nucleotide Archive accession PRJEB43881. MDT was additionally supported by ERC investigator grant no. 787282-B2C.

## Authors’ contribution

CR, KGD, and MM designed study; CR, KGD, VM, AH, and FM performed laboratory work; CR and MDT performed bioinformatic work; GRP was responsible for SEM analysis; HD, HL, ZL, RK, HF, AA, TS, and MM worked directly with and provided archaeological samples. All authors contributed to writing the manuscript, approve this study, and are accountable for all aspects of the work.

## Acknowledgements

We thank Daniel Bradley for his mentorship, discussions and reading of the paper; We thank Andreas Steiger-Thirsfeld for SEM images and N. Tehrani for sample photography.

