## Supplementary Material for "Exceptional ancient DNA preservation and fibre remains of a Sasanian saltmine sheep mummy in Chehrābād, Iran"

### Contextual Information for Archaeological Sites

#### Chehrābād, Iran

The mummified sheep leg 4305 (Supplementary Figure 1) was discovered during Iranian-German research work in one of the debris layers (feature 31091) in the northwestern edge of the mine during modern archaeological excavations of the Douzlākh saltmine of Chehrābād [1]. At the northwestern edge, part of a large mining gallery was discovered. This part had collapsed between 405 and 380 BCE but was reopened in the Early Sasanian period ($2^{nd}$-$3^{rd}$ centuries CE). The working area in the northwestern edge was partly refilled most likely between the $4^{th}$ and $5^{th}$ century CE, of which the layer 31091 belongs. The dating of the sheep-leg (Table 1, Supplementary Figure 3) is concurrent to the layer dating (reference [1], 43-45, Table 11). It was most likely deposited in the course of refilling activities during a salt extraction phase when mining took place on the northwestern salt rock edge of the mine.

A skin cutting of sheep leg 4305, MUM2 (Supplementary Figure 2), was used for DNA extraction and $C_{14}$ date estimation.

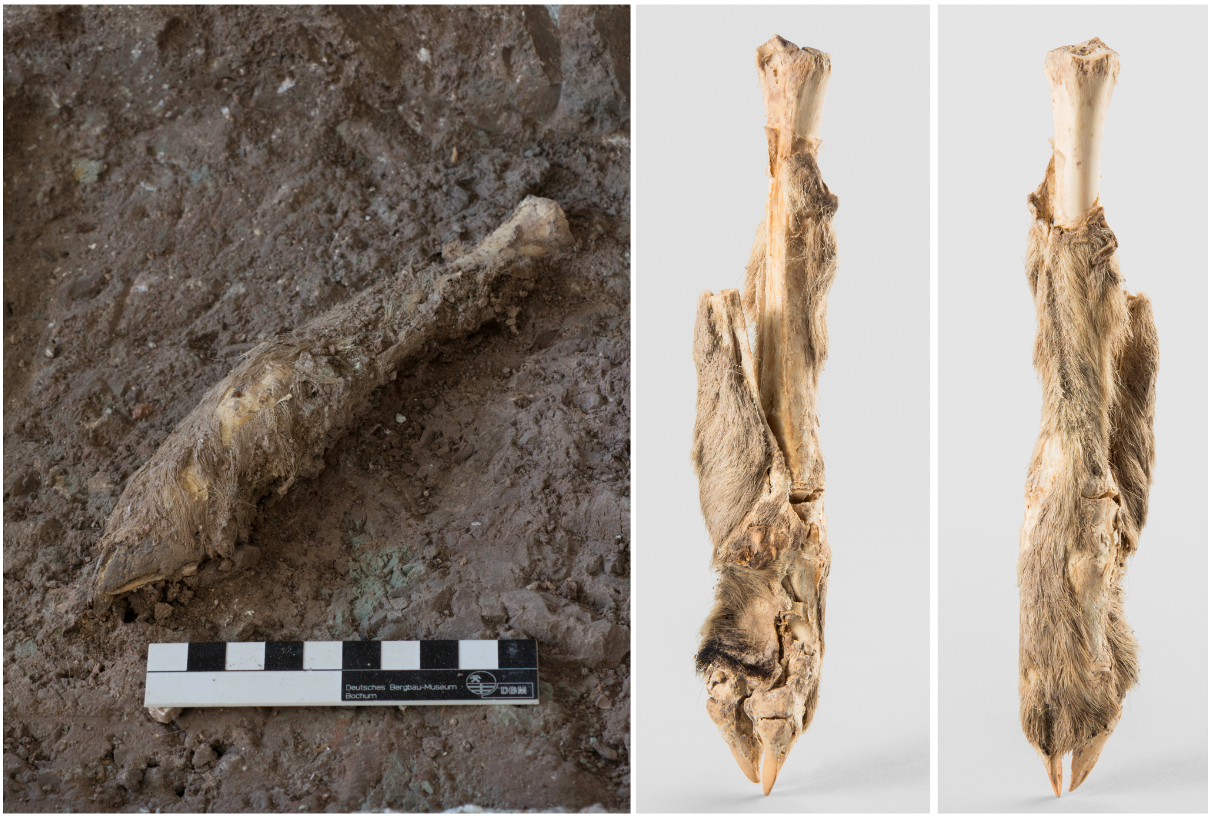


**Supplementary Figure 1:** Sheep leg (4305) in the archaeological context of layer 31091 (left) and recent condition after cleaning (two angles, centre, right), Photos: DBM/RUB/AMZ, Thomas Stöllner (left), N. Tehrani (centre, right).

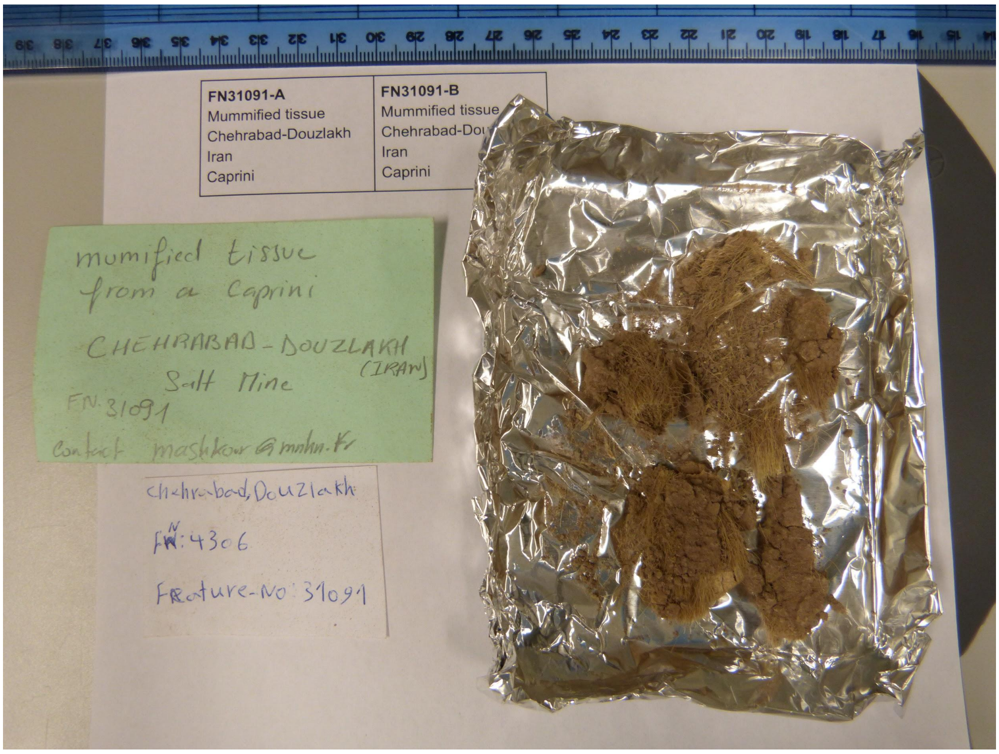

**Supplementary Figure 2. Photograph of the mummified skin of MUM2.** MUM2 was sampled from the sheep leg 4305.

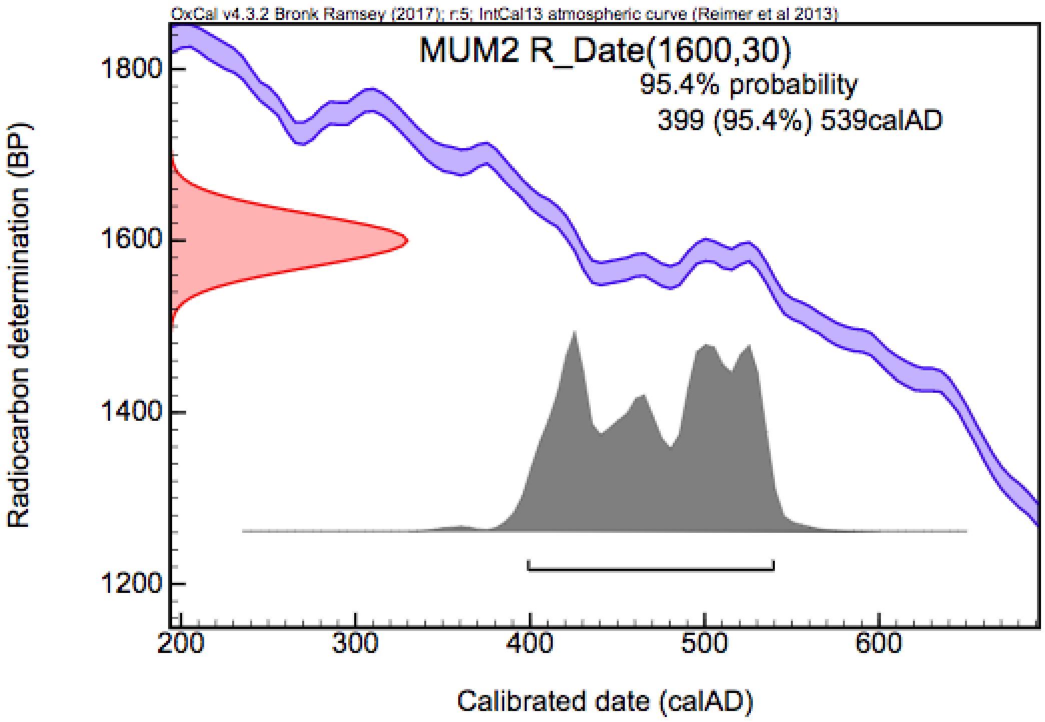


**Supplementary Figure 3: OxCal 4.3.2** [2] **calibration of the radiocarbon age of MUM2 using IntCal13** [3].

### Tepe Hasanlu, Western Azerbaijan, Iran

Tepe Hasanlu is one of the key sites of northwestern Iran, due to its long-term occupation and

well-defined stratigraphy. It is located in the Solduz valley on the southern shore of Lake Urmia, Western Azerbaijan province of Iran (1043 m ASL, Latitude: 37°0'16.15"N, Longitude: 45°27'31.74"E). Robert H. Dyson Jr. directed ten seasons of excavations at Hasanlu from 1956 to 1977 [4,5]. The site was occupied during ten cultural periods from the Late Neolithic (period X) to the Ilkhanid dynasty (period I) [6]. The most represented periods in the site are the Late Bronze (period V) and Iron Age (period IV-III). The Iron Age II citadel of Hasanlu was destroyed and fired during a battle, probably by the Urartian army, around 800 BC [7,8]. Hasanlu period IIIc and b, are attributed to Iron Age III (Urartian period / 800-600 BCE) and period IIIa allocated to the Achaemenid Empire (550-530 BC). The citadel with fortification wall, a few massive public buildings like garrison quarters, flimsy structures, stables and other architectural remains had been discovered from period III [9]. Extensive excavations in these periods resulted in the discovery of burned buildings, thousands of artifacts in closed contexts, and a large quantity of human and animal bone remains. Animal bones are among the most abundant material recovered in the site. Despite the taxonomic diversity of the remains in Hasanlu, domestic sheep, goat, and cattle were the most exploited animals in all the periods and cattle contributed much less to the diet than sheep and goat during the Bronze Age and Iron I to II, while its contribution increased to as much as Caprini during Iron III and Historical periods [10,11]. The sheep sample from Tepe Hasanlu analysed in this study was a petrous bone recovered from deposits of level IIIb (Iron Age III) in campaign 1974, Op. X32, Stratum 5, Lot 26.

**Azer2 Azer2 MM TH3**

### Nishapur, Khorasan, Iran

Nishapur, one of the major urban centers of Khorasan, is located in Northeast Iran on a plain limited by the Binalud heights in the North and the Kashmar heights in the South (1198 m ASL, Latitude: 36°10'16.72"N, Longitude: 58°50'50.82"E). Nishapur as a major social and economic urban area exists in a cultural landscape in which the economic life of rural and urban entities was based on agriculture, horticulture, and animal husbandry. Nishapur had been an important producing center of silk, wool, cotton and textiles and their export throughout the Islamic world and beyond [12]. Archaeological research in Nishapur began by the Metropolitan Museum of Art during the 1930s [13–17]. Since then, surveys and excavations have been realized by national and international missions. Excavations conducted by the Iranian-French team resulted in the discovery of a significant amount of animal remains in the citadel [18,19]. Recently, a new Iranian excavation that has been carried out in the citadel and the urban area also produced substantial bioarchaeological data [20]. 2470 pieces of animal bones from the first collection found in 2005-2006 were studied in the Archaeozoology section in the Bioarchaeology Laboratory of the Central laboratory of the University of Tehran [21]. The remains could be dated from the Sasanian period to the 12th century CE. The results indicate that sheep, goats, and cattle were the most exploited animals for both meat and secondary products. The Nishapur petrous bone sample analyzed in the present study was discovered in the campaign 2006, Tr. 15B located in the north part of the citadel, Square: JXV, 6-12th century CE.

**Khor1 Khor1 MM NKD1**

### Supplementary Methods

#### DNA extraction

Sample preparation, extraction and library preparation were performed in a dedicated ancient DNA laboratory in the Smurfit Institute of Genetics, Trinity College Dublin. A small section of skin (0.454g) was cut using a disposable blade. Prior to the extraction the sample was washed with 1ml of water (lab grade and UVed) then with 600ul of Ethanol (99%) and then again with 1 ml of water. Each time the supernatant was removed after centrifugation. DNA was extracted using a modified version of the classic phenol/chloroform DNA extraction protocol [22]. A modified extraction buffer was used, where 3mM CaCl2 and 30mM DTT were added to the original extraction buffer.

#### Library preparation and sequencing

Treatment of ancient DNA with Uracil-DNA glycosylase (UDG) and Endonuclease VIII has been shown to remove base misincorporations caused by deamination in aDNA [23]. For UDG-treated libraries, 5 µl USER® Enzyme (1,000U/ml; Uracil-Specific Excision Reagent, New England BioLabs Inc.) was added to 16.25µl purified DNA and incubated in an Eppendorf ThermoMixer for 3 hours at 37°C, prior to library preparation.

Both UDG-treated and not-treated libraries were constructed based on the protocol of [24] with modifications as reported in [25]. Control tubes (21.25µl $H_2O$ starting material) were carried out for library construction. Indexing Polymerase Chain Reactions (PCRs) were performed using Accuprime *Pfx* Supermix (Invitrogen), primer IS4 (10 µM) and a unique indexing primer (5 µM) as detailed in [24]). The concentration of amplified libraries was quantified with a TapeStation 2200 (Agilent).

Three rounds of sequencing were done: single-end and paired-end shotgun sequencing of USER-treated libraries on Illumina HiSeq 2500 (Macrogen), and single-end shotgun sequencing for non USER-treated libraries on Illumina MiSeq (TrinSeq). The UDG-treated library was sequenced on a HiSeq 2500 Illumina platform (100bp SE and 100bp PE) via a commercial sequencing company (Macrogen, Republic of Korea) while the non-UDG-treated library was sequenced on a MiSeq Illumina platform (50bp SE) at TrinSeq (Trinity College Dublin, Ireland) with Phi X control at 1%.

#### Raw-read processing

The quality of fastq files was assessed using FastQC [26]. For single-end libraries, adapters were trimmed from reads of raw fastq files using cutadapt 1.9.1 [27] (cutadapt -a AGATCGGAAGAGCACACGTCTGAACTCCAGTCAC -O 1 -m 30). This removed reads shorter than 30bp and trimmed adapter sequence if an overlap of more than 1 bp was found between the read and adapter sequence.

For paired-end libraries, adapters were trimmed using AdapterRemoval v2.1.1 [28] (AdapterRemoval --file1 reads_1.fq --file2 reads_2.fq --basename output_paired --trimns --trimqualities --minquality 25 --collapse). The --trimns --trimqualities --minquality flags cause stretches of consecutive Ns and/or bases of quality lower than 25 from the 5' and 3' termini to be trimmed. The --collapse flag merges mate pairs that overlap by at least 11 bp into a single read. Base quality scores of collapsed reads are then recalculated using the metadata of the paired reads. The collapsed fastq file and the truncated read pairs were used in downstream analysis.

### Read alignment

All trimmed reads of samples were aligned to OviAri3.1 using BWA [29] 0.7.5a with relaxed parameters (-l 1024 -n 0.01 -o 2). These flags disable seeding, allow for more mismatches with the reference and allow 2 gap openings, respectively.

For single-end reads, the resulting sai files were converted to sam file format using BWA samse [29]. Paired-end reads were handled using BWA sampe [29] to collapse sai pairs into one sam file. Sam files were then converted into the binary bam files using SAMtools (v1.7) view [30]. The -F4 flag was used with SAMtools view to discard unmapped reads for all bam files. For paired-end reads, the -f2 flag was also used to remove pairs of reads where one paired read was aligned to a different chromosome. The bam files were then sorted using SAMtools sort and PCR optical duplicates were removed using SAMtools rmdup. Bam files were then filtered for reads of mapping quality less than 30 using SAMtools view. Read groups were added to each bam file using Picard AddOrReplaceReadGroups (https://broadinstitute.github.io/picard/). Finally, these filtered bam files were merged using SAMtools merge. Genome coverage was calculated using GATK DepthOfCoverage [31]. Alignment statistics are summarised in Supplementary Table 1.

| Sample | Library type | Sequencing platform | Raw Reads | Trimmed Reads | Aligned Reads | MQ30, rmdup Aligned Reads | Endog. % |
|---|---|---|---|---|---|---|---|
| **MUM2** | Non-USER treated | Illumina MiSeq 63 SE | 6,371,514 | 6,317,966 | 2,479,430 | 1,959,646 | 31.02 |
| **MUM2** | USER treated | Illumina HiSeq 100 SE | 63,389,209 | 62,636,422 | 19,172,915 | 15,576,961 | 24.87 |
| **MUM2** | USER treated | Illumina HiSeq 100 PE | 598,909,352 | 348,606,473 | 109,018,213 | 86,945,474 | 24.94 |
| **Khor1** | Non-USER treated | Illumina MiSeq 65 SE | 333,116 | 319,697 | 263,477 | 186,846 | 58.44 |
| **Khor1** | USER treated | Illumina HiSeq 100 PE | 6,230,996 | 3,146,933 | 2,796,988 | 1,934,890 | 61.48 |
| **Azer2** | Non-USER treated | Illumina MiSeq 65 SE | 1,430,246 | 1,379,462 | 645,402 | 432,134 | 31.33 |
| **Azer2** | USER treated | Illumina HiSeq 100 BP | 20,448,674 | 10,835,702 | 5,029,767 | 3,074,445 | 28.37 |

**Supplementary Table 1: Sequencing and alignment statistics.**

### Damage patterns

Bam files were assessed using mapDamage2.0 [32]. mapDamage2.0 calculates read length distributions and substitution patterns at the 5' and 3' ends of read fragments of aligned bam files. mapDamage2.0 also outputs a number of damage parameters estimated in a Bayesian manner including δS, the cytosine deamination probability in single strand context. Published sequencing data of other ancient samples were retrieved from NCBI GenBank in order to contextualise the damage patterns of our sample with published data (Supplementary Table 3).

Depurination levels were inferred using frequencies of purines close to the 5' end of fragment strand-breaks. The use of *T4* DNA polymerase in our library preparation removes 3' overhangs, therefore only 5' ends are considered. Only non-UDG libraries were used to calculate these levels as USER excises uracil residues in DNA fragments, creating new strand breaks. Using Qualimap v2.1.3 [33], nucleotide frequencies were calculated on filtered bam files at each position of DNA fragments (Supplementary Figure 4). Following the methods detailed in [34], approximate depurination rates were calculated.

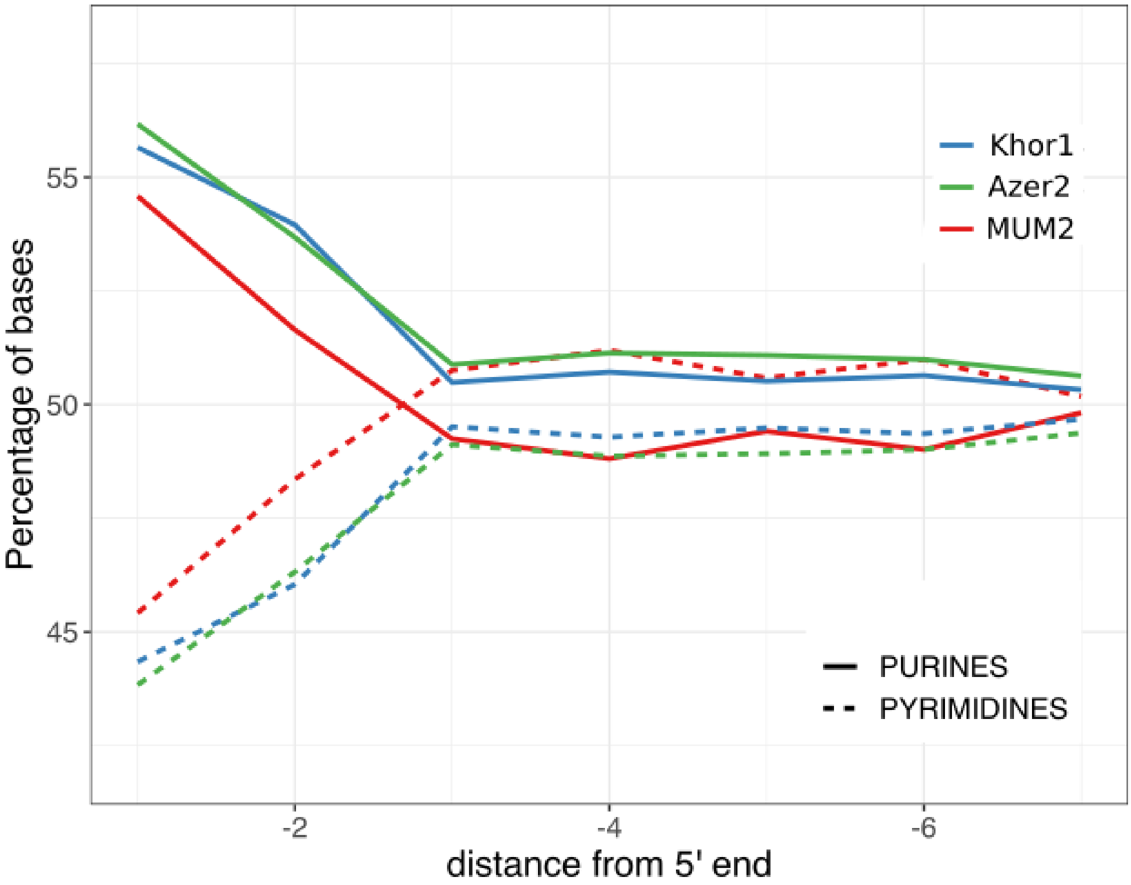


**Supplementary Figure 4: Nucleotide frequencies at 5' end of DNA fragments.** The depurination rate was 6.3%, 5.6% and 5.3% for MUM2, Khor1, Azer2, respectively.

### Metagenomics

For all metagenomic analysis, we first removed reads originating from the host organism and human contamination. To do this, all trimmed reads were aligned to the sheep genome (oviAri3.1) and the human genome (hg38) using methods detailed above. Unmapped reads were retained using SAMtools view -f 4 and converted to fastq files using SAMtools fastq. The retained reads were then deduplicated using PrinSeq [35]. In total 202,815,624 reads were retained for MUM2 (176,858,319 collapsed/single end reads and 25,957,305 paired end reads). This was repeated for Khor1 and Azer2, resulting in 281,649 and 4,315,523 reads, respectively.

Unmapped, deduplicated reads were searched against the Kraken2's standard database using default settings [36]. This database includes sequences of bacterial, archaeal and viral genomes. Kraken2 classified 11.96% (24,247,293 reads) of the unmapped, deduplicated reads using this database. To calculate Class abundances, microbial abundances were recalculated using Bracken [37] est_abundance.py, with a minimum threshold of 0.1% of total classified reads. 57.13% (13,499,165) of the classified reads belonged to the salt-adapted *Halobacteria* Class which reflects the high salinity environment of Chehrābād.

| Class | Total Reads Assigned | % of Assigned Reads | Domain |
|---|---|---|---|
| *Halobacteria* | 13499165 | 57.13 | Archaea |
| *Actinobacteria* | 3879524 | 16.42 | Bacteria |
| *Clostridia* | 1243905 | 5.26 | Bacteria |
| *Alphaproteobacteria* | 914412 | 3.87 | Bacteria |
| *Gammaproteobacteria* | 823160 | 3.48 | Bacteria |
| *Betaproteobacteria* | 659427 | 2.79 | Bacteria |
| *Bacilli* | 295377 | 1.25 | Bacteria |
| *Other and Host* | 155083 | 8.54 | N/A |

**Supplementary Table 2: Taxonomically assigned reads of MUM2 by Kraken2, recalculated by Bracken, at Class level.** The salt-adapted Class *Halobacteria* predominates with 57.13% of assigned reads.

Using the same methods, *Halobacteria* were found in very low abundance in our comparative bone samples, Khor1 and Azer2 (0.7% and 0.67% of assigned reads, respectively with a threshold of 0.1%). As a comparison with other ancient skins, metagenomic analysis of the Tyrolean Iceman's preserved clothes [38] and ancient parchment [39] also revealed very low levels of *Halobacteria* present (0-0.02% and 0-0.3% of assigned reads, respectively). The uniquely high levels of halophilic archaea present in MUM2 is suggestive of the large effect the saltmine environment had on the microbial communities.

We used SourceTracker2 to further define the metagenomic composition of MUM2. SourceTracker2 is a Bayesian source-prediction tool that can estimate the proportion of sequencing reads of samples (sinks) originating from selected metagenomes (sources) [40]. DNA sequences of various relevant metagenomes (salt-rich, live sheep skin, soil, tissue decomposers, and laboratory reagents) were downloaded from NCBI (Supplementary Table 4). The sequences were trimmed using

AdapterRemoval v2.1.1 and taxonomic assignment for these reference sequences was done by Kraken2, as detailed above. The kraken reports were merged and converted to a BIOM table using kraken-biom (https://github.com/smdabdoub/kraken-biom) and the human taxid (9606) was removed.

Source prediction of MUM2, Azer2 and Khor1 was done at both Species and Genus level using default settings. We were unable to use the “Laboratory reagent” source at Genus level as there was an insufficient number of taxonomically assigned Genuses (27) for our rarefaction level (1,000). At Species level, SourceTracker2 estimates 0.4725 of the microbial community of MUM2 originates from a salt-rich environment., while only 0.003-0.007 of the comparative bone samples were estimated to be from this environment (Table 2; Supplementary Figure 5A). This estimation of salt rich increases to 0.746 when estimated at Genus level (Table 2; Supplementary Figure 5B).

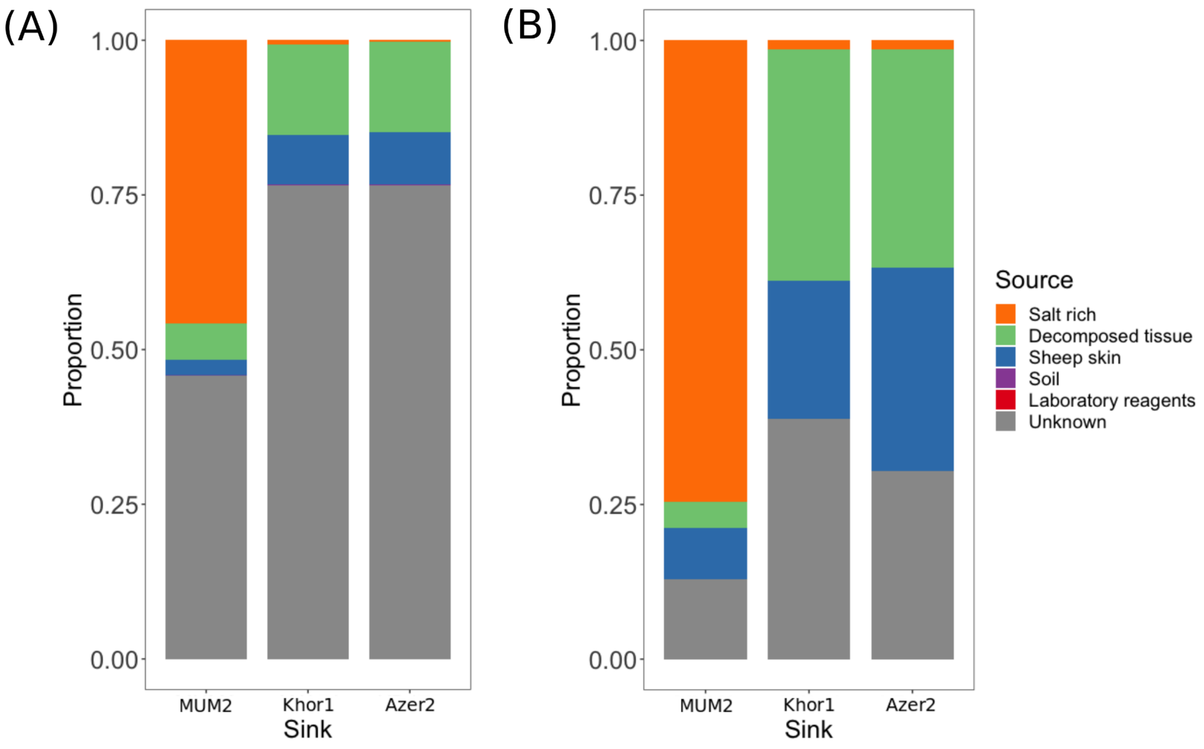


**Supplementary Figure 5: Prediction of metagenomic source proportions by SourceTracker2 at (A) Species level and (B) Genus level.**

In order to cross-validate our results, and to investigate individual bacterial species, MIDAS [41] was selected as it has been reported to be the amongst most accurate for aDNA microbial communities [42]. This tool quantifies bacterial species abundance from shotgun metagenomes. MIDAS clusters more than 31,000 bacterial reference genomes into nearly 6,000 species groups based on sequence identity of 30 universal marker genes.

We observe a large proportion of detected microbial species in MUM2 (8 of the 9 most abundant species) as being salt adapted or salt tolerant (Supplementary Table 4). Of these, just one species is identified in ancient sample Azer2. This species, *Streptomyces sp 59965*, is a generalist. Similarly, none of the defined salt adapted bacteria in MUM2 were detected in shotgun data of leather from the Tyrolean Iceman ([38], results not shown). Top MIDAS hits for ancient parchment shotgun data ([39], results not shown) largely comprised of bacteria associated with animal skins (e.g. *Erysipelothrix rhusiopathiae*, *Propionibacterium acnes*, *Staphylococcus equorum, Brevibacterium linens*). Some salt-tolerant species (Table 2) were found in low abundance (<2%) of several ancient parchment shotgun data except for *Saccharomonospora glauca 62535* which was found at 4% abundance in sample SAMEA104143134. Overall, these results support the observation that MUM2's microbial profile reflects its unique taphonomic context.
To validate the authenticity of the most abundant bacterium detected, all sequencing data was aligned to GCF_00371785.1, an *Actinopolyspora halophila* reference genome (Actinopolyspora halophila DSM 43834, retrieved from NCBI GenBank). A 0.33X genome was generated (std per bp = 1.13X). Using Qualimap v2.1.3 [33], the coverage of this bam file was visualised. The coverage of the bam file was even, as is recommended as a validation in [43]. The median read length of the aligned reads was 74 bp.

### Controls

Controls that were done throughout the sample preparation, extraction and library preparation were also screened for halophilic signatures. The number of reads taxonomically assigned by Kraken2 was low (643 - 4648 assignments) and in all the presence of the *Halobacteria* clade was low (0.009-0.01). Metagenomic source prediction was done at a Species level using the same SourceTracker2 dataset as above (Supplementary Figure 6). The salt-rich environment was not predicted to contribute highly to any of the controls (0.003-0.024). After “Unknown”, “Laboratory reagents” were predicted to be the highest contribution to all controls except the Library Control. Here, SourceTracker2 predicted 0.66 of its metagenome to belong to “Sheep skin”, suggesting possible skin contamination during library preparation.

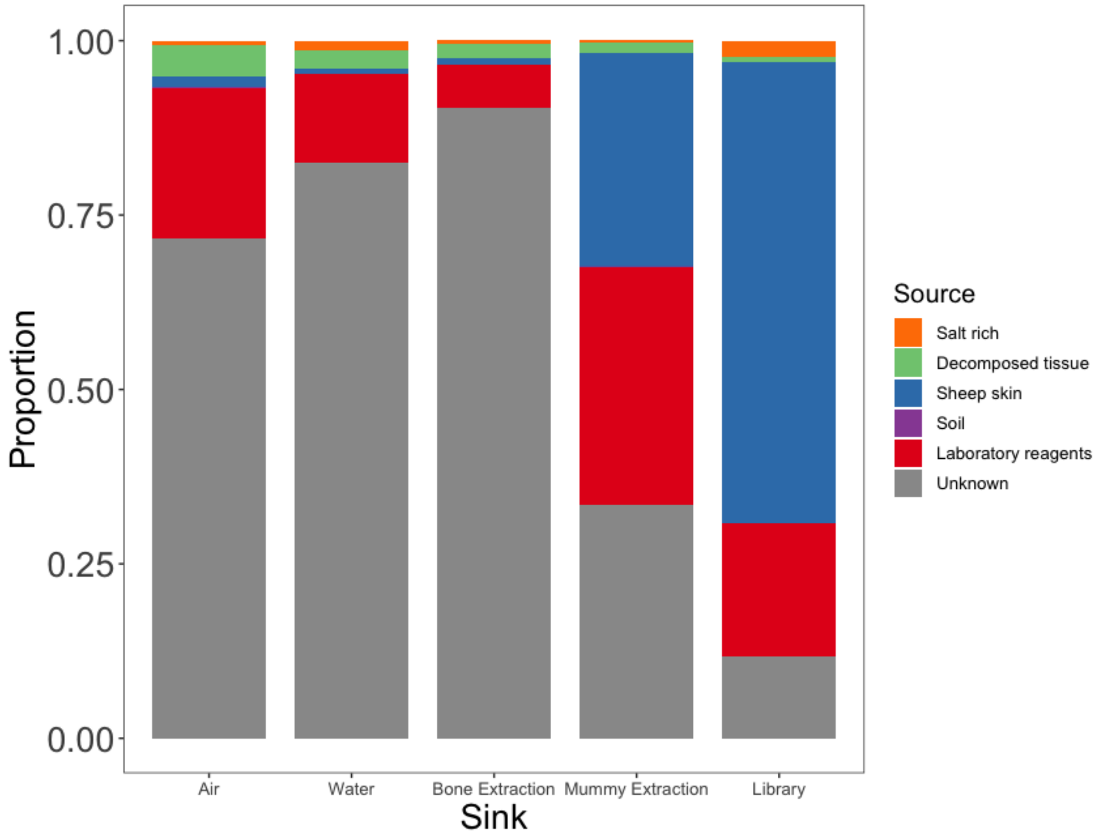


**Supplementary Figure 6: Prediction of metagenomic source proportions of controls by SourceTracker2 at Species level.**

### Mitochondrial tree

All UDG-treated sequencing data was aligned to the domestic sheep mitochondrial reference, AF010406.1 (retrieved from NCBI GenBank) using an identical method as to the nuclear genome alignment. Consensus fasta sequences were generated using ANGSD [44] (angsd -doFasta 2 -doCounts 1 -setMinDepth 3 -minQ 20 -minMapQ 30). With these flags, base calls must have a minimum quality of 20 and a minimum MAQ of 30 to be considered. In the event that there are more than one base at a given site, the most common base is chosen. For a site to be called a minimum depth of 3 is required.

Ten modern sheep whole genome sequences and one urial sheep whole genome sequence published in [45] were retrieved from NCBI GenBank. Multiple sequence alignments were performed with MUSCLE [46]. A maximum-likelihood phylogeny (100 bootstraps) was generated. The HKY85 substitution model was selected for tree-building using jmodeltest2 [47,48]. Alignments and maximum-likelihood phylogeny constructions were implemented in SeaView v. 5.0.1 [46,49,50].

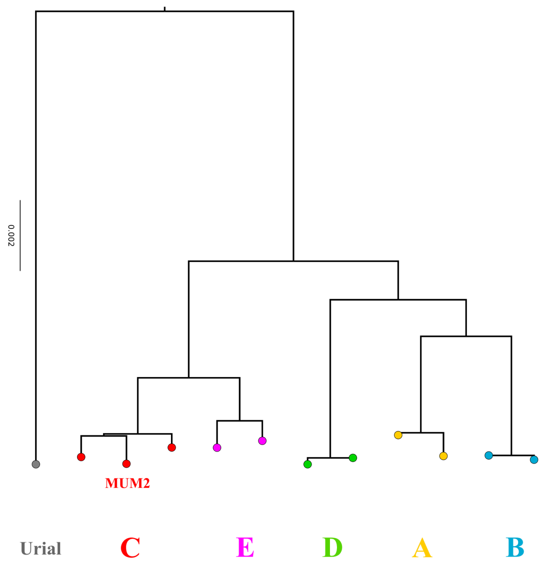


**Supplementary Figure 7: Whole mitochondrion maximum-likelihood phylogeny of domestic sheep (100 bootstraps).** MUM2 falls within Clade C. All clades of the phylogenetic tree are well-supported by bootstrap values (>0.97).

### SNP Calling

The ovine SNP50 HapMap dataset used for the analyses described was provided by the International Sheep Genomics Consortium and obtained from www.sheephapmap.org in agreement with the ISGC Terms of Access. This dataset is genotyped on the Illumina OvineSNP50 Genotyping BeadChip (Illumina), covering >54,000 SNPs evenly spaced over the sheep genome. 1503 genotyped individuals are included in this dataset, representing modern breeds from Africa, the Americas, Europe, SW Asia and Asia [51].The genomic positions of the OvineSNP50 were updated for the OviAri3.1 genome build (https://doi.org/10.6084/m9.figshare.8424935.v2) and SNPs flipped to the forward strand. Using PLINK v1.9 [52], these SNPs were filtered down to 44,223 sites using a MAF filter of ≤ 0.5.

The coverage of our sample was too low (approx. 4X) for accurate diploid genotype calling. To circumvent this issue, the pileup tool in GATK [31] v3.7 was used to report sample base calls for each read at each position of the filtered SNP list (44,223 SNP positions). A minimum base quality of 30 was required to be considered and sites with three or more different bases present were removed. A single base call was then chosen at random from each site and duplicated to create a pseudohaploid homozygous genotype at that position for each individual. In total 43,027 SNPs were called for MUM2.

Using PLINK v1.90, the pseudohaploidised MUM2 genotype calls were merged with the modern sheep breeds dataset. In cases where the allele of MUM2 matched neither of the alleles given in the sheep breed position, the SNP position was flipped using PLINK v1.90 and attempted to be merged again. Any remaining triallelic SNPs were removed. The final merged dataset was used for *f3* statistics, ADMIXTURE and TreeMix analysis.

### PCA

Projection PCA using Procrustes analysis was performed using LASER v2.03 [53]. The PCA reference space and projection transformation were constructed using a subsample of our modern breeds dataset. This subsample contained representation from all modern breeds, chosen randomly. Using the OvineSNP50 reference panel, pileup files were generated for MUM2, Khor1 and Azer2 using SAMtools mpileup (-q 30 -Q 20). Each sample was projected onto this PCA space and averaged across ten independent runs. The uncertainty based on the standard deviation * 2 for each sample was plotted on the map (Supplementary Figure 8).

The plot of PC1 vs PC2 shows that PC1 differentiates modern European breeds from breeds of the rest of the world. PC2 differentiates non-European breeds. A cline from African, Asian and South West Asian appears. These breeds do not create discrete clusters, however, and some individuals from different locations can be seen to cluster together in geographically-heterogeneous groups. MUM2 falls with the main cluster of SW Asian breeds. Khor1 and Azer2 both project within the diversity of SW Asian breeds but fall away from this main cluster on PC1. However we note that the uncertainty of PC1 of Khor1 and Azer2 is large, likely due to the lower SNPs called while the coordinates of MUM2 had a low uncertainty.

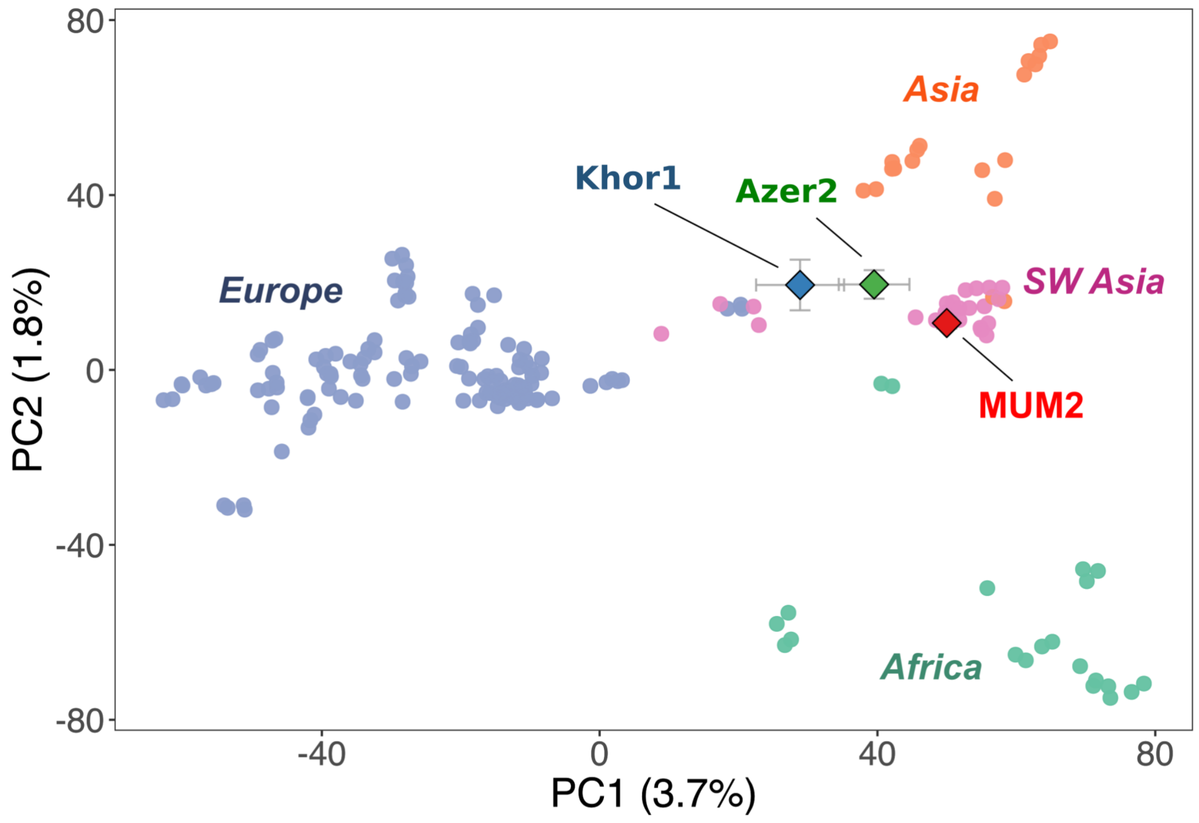


**Supplementary Figure 8: PCA of global sheep affinity and Procrustes projection of MUM2, Khor1 and Azer2.** Error bars represent the standard deviation of each PC coordinate between independent runs * 2. PC1 separates breeds of

European origin and the rest of the world, while PC2 separates Asian and African breeds into an approximate cline of Central/East Asian, SW Asian, African [51].

### Local affinity

To investigate local affinity, a PCA was produced using only breeds of SW Asia and Asia using LASER v2.03. Here, PC1 differentiates Asian and SW Asian breeds. PC2 then separates one SW Asian (Turkish) breed from all other SW Asian breeds. When MUM2, Khor1 were projected onto this reference space using LASER, they clustered with SW Asian breeds rather than those from SE Asia (Supplementary Figure 9). Azer2 falls away from this cluster of SW Asian breeds, however, there is a larger amount of uncertainty again due to lower SNPs called.

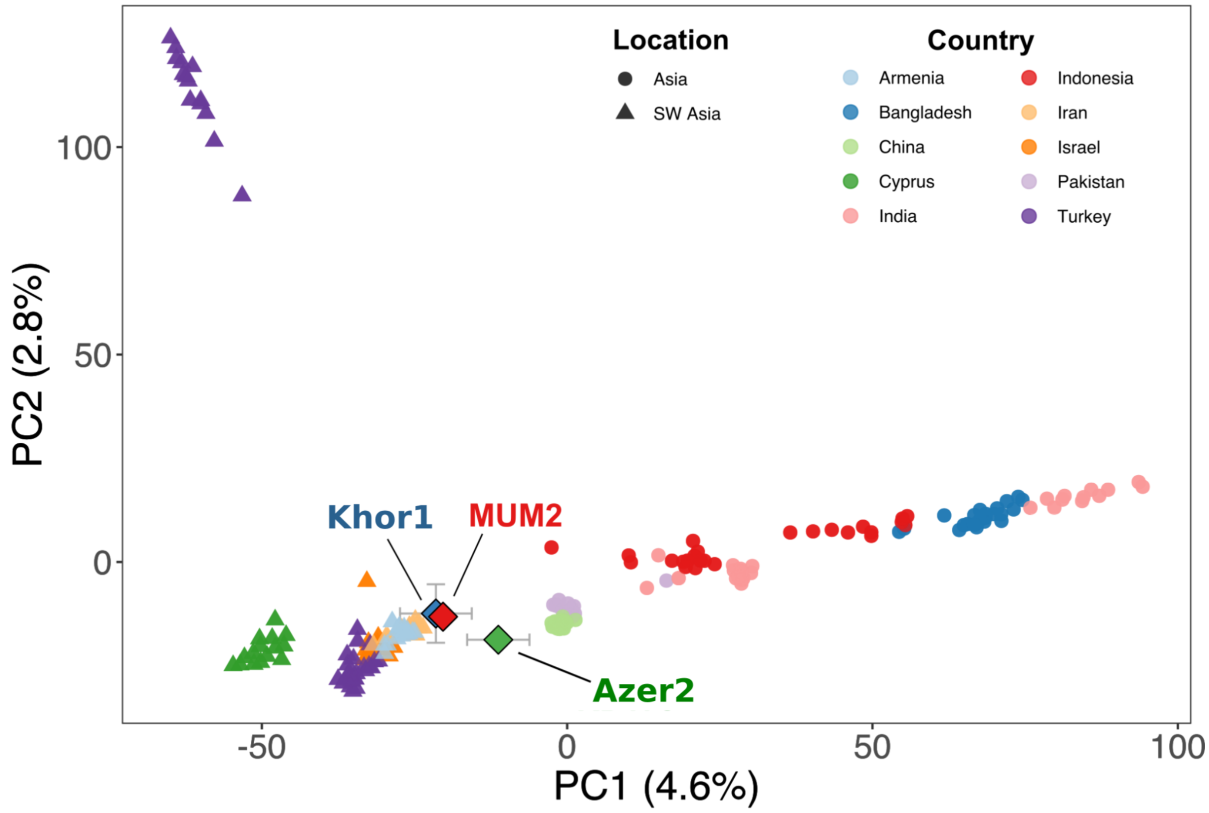


**Supplementary Figure 9: PCA containing only Asian breeds.** Error bars represent the standard deviation of each PC coordinate between independent runs * 2. MUM2 clusters with Southwest Asian breeds. Some amount of genetic flux in local breeds in the past 1600 years is inferred by MUM2 falling slightly away from the main cluster of Southwest Asian breeds. Azer2 falls further away from the main cluster of

SW Asian breeds, however, we note increased relative uncertainty of its PC1 position due to the lower number of called SNPs.

### $f_3$ Statistics

$f_3$ statistics were computed to quantify the amount of genetic drift shared between MUM2 and modern sheep breeds. Three Asiatic mouflon individuals that were genotyped on the OvineSNP50 Genotyping BeadChip were used as an outgroup. Using PLINK v1.9, these were merged with our full dataset of modern breeds and MUM2. Using PLINK v1.9, a filter of 0 for missingness was applied to this merged dataset. The dataset was then filtered for linkage disequilibrium (LD) between markers, with the command --indep-pairwise 50 5 0.5. Outgroup $f_3$ values were not estimated for Khor1 and Azer2 due to the lower number of called SNPs.

Using this complete dataset, we calculated the $f_3$ statistics in the form of $f_3$(MUM2, modern breed; Asiatic Mouflon) with ADMIXTOOLS [54]. $f_3$ statistics were plotted using approximate geographic origins of breeds with R using the sf package [55] (Figure 2A).

### TreeMix

The complete dataset used was converted into TreeMix format with the plink2treemix.py utility [56]. TreeMix [56] was used to construct a model of population splits with no migration events. The tree was rooted with Asiatic mouflon and blocks of 500 SNPs were selected (-k 500). No sample size correction was used (-noss). Clades emerge based on the geographic location of the sheep breeds. MUM2 falls in the SW Asian clade and forms a branch with the breed sampled from Iran, Afec-Assaf.

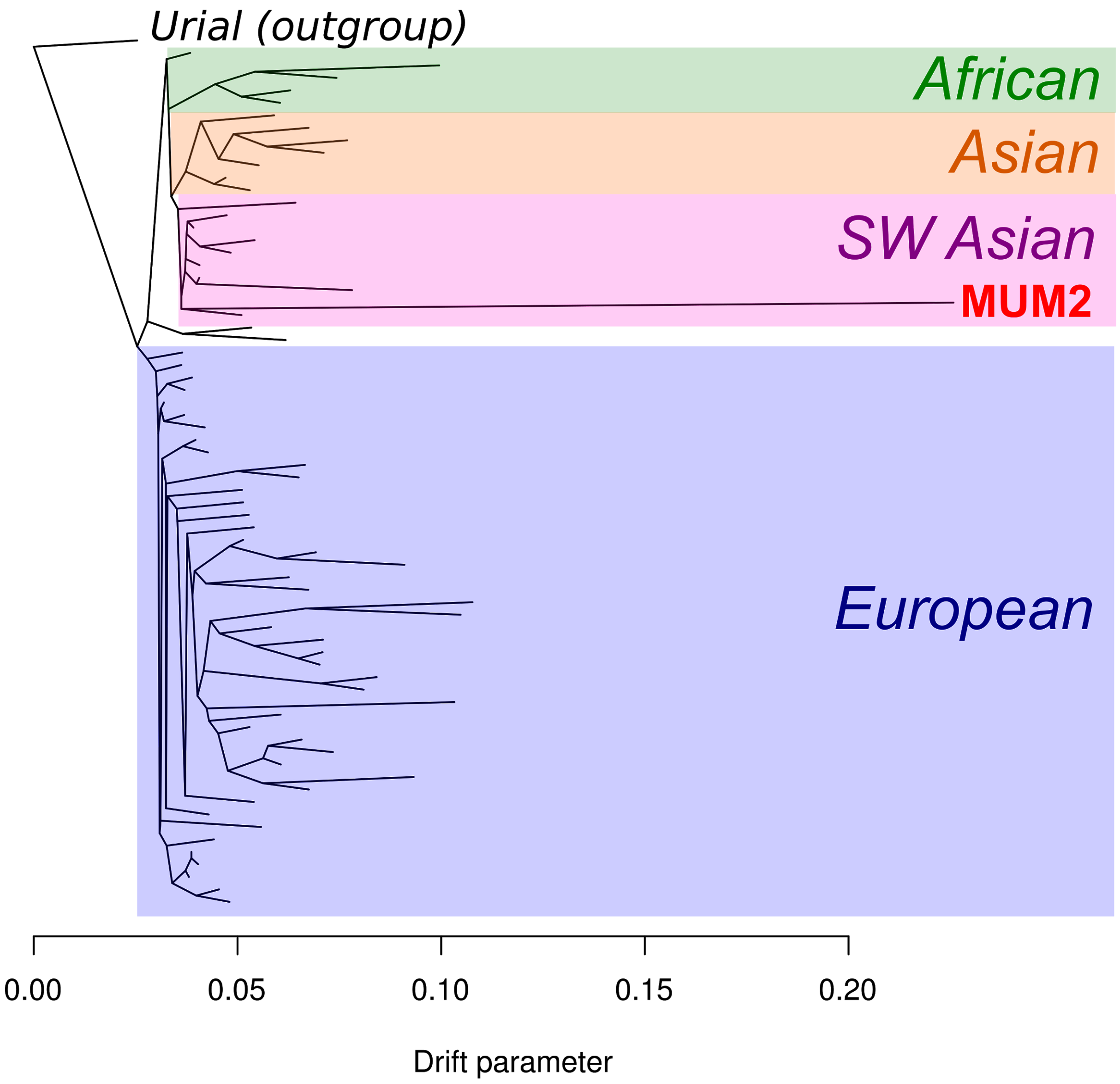


**Supplementary Figure 10: TreeMix plot with 0 migration edges.** MUM2 falls into the SW Asian breed clade. MUM2 forms a branch with the modern Iranian Afec-Assaf breed.

### ADMIXTURE

The genotyped breed dataset was downsampled using PLINK v1.9 to include approximately 90 individuals from each Asia, Europe and Africa. Using PLINK v1.9, a filter of 0 for missingness was applied to this merged dataset. The dataset was then filtered for linkage disequilibrium (LD) between markers, with the command --indep-pairwise 50 5 0.5. Finally, transitions were removed using PLINK v1.9 and VCFtools [57]. This fully filtered dataset contained 31,261 sites.

Unsupervised ADMIXTURE was explored with a range of hypothetical ancestral populations, *K*s (2-20). ADMIXTURE runs were replicated three times for each K and set a different random seed for each. The three replicated runs for each K were averaged, and the cross-validation values which describe the approximate error of each ADMIXTURE run were inspected [58]. The run with the lowest cross-validation error was selected for presentation, *K*=13. In this ADMIXTURE analysis, the predicted ancestral profile of MUM2 most resembles Qezel, a sheep breed local to Iran.

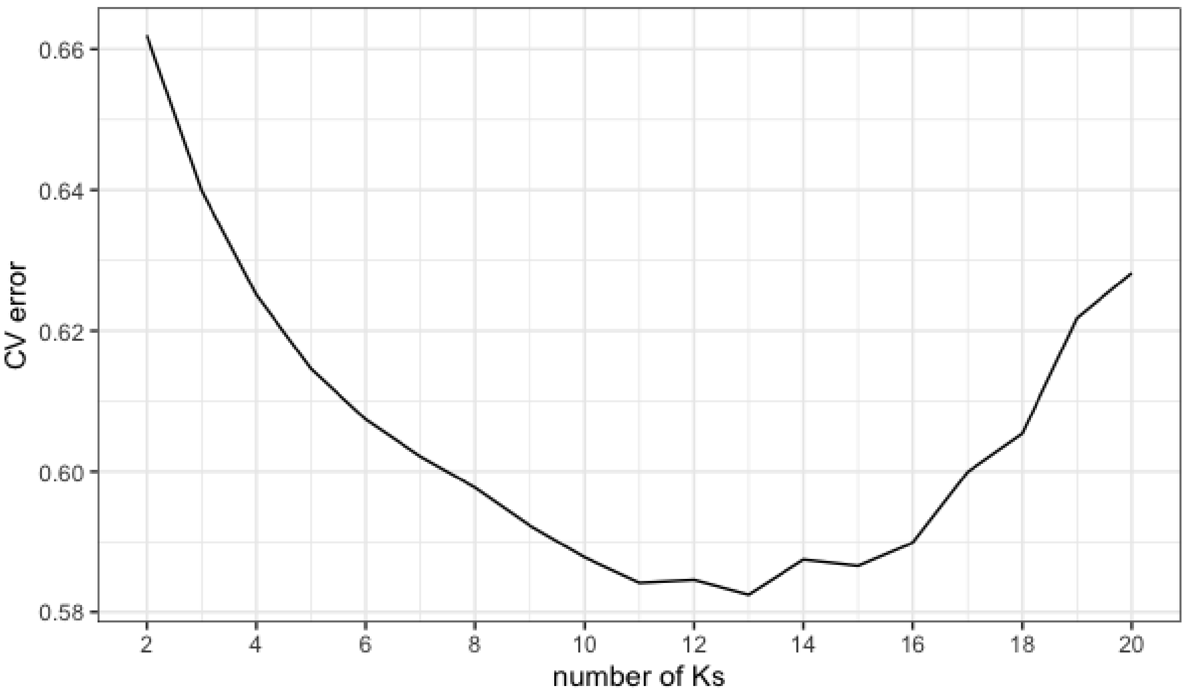


**Supplementary Figure 11: Average cross-validation values of unsupervised ADMIXTURE for each *K*, hypothetical ancestral population.**

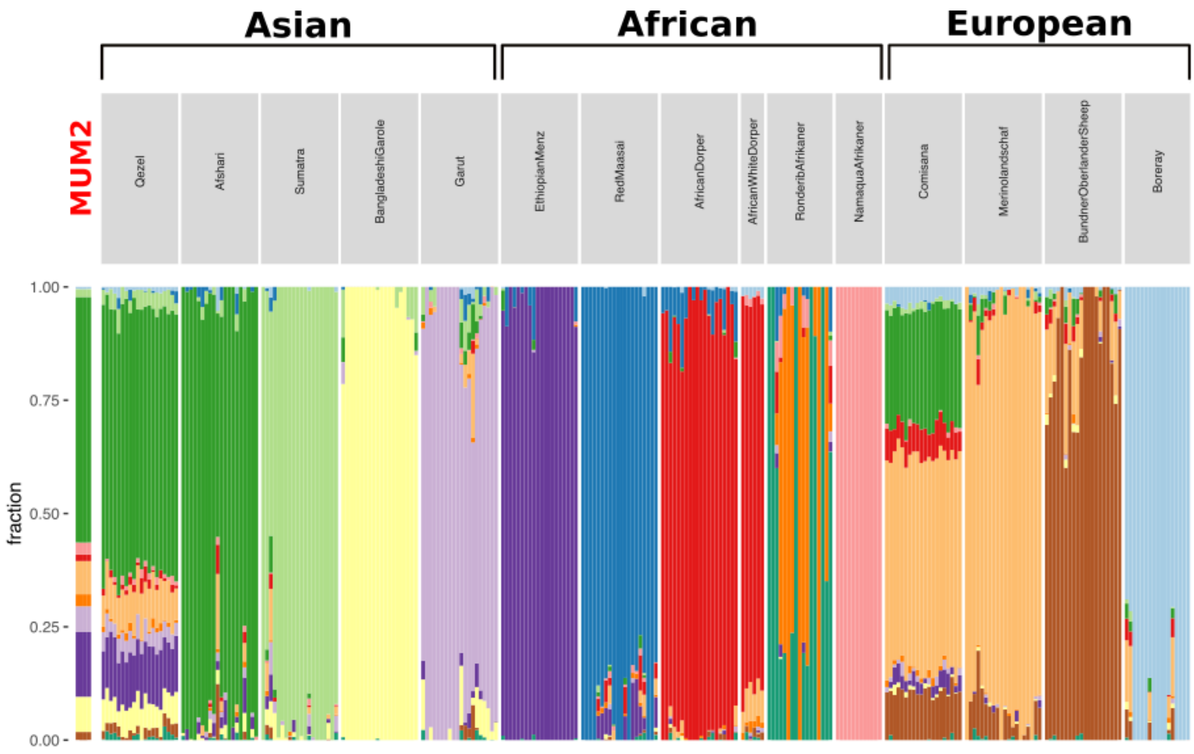


**Supplementary Figure 12: ADMIXTURE plot with K=13 ancestral components.** MUM2 displays a similar ancestral profile to modern Iranian sheep, represented by the Qezel breed.

### Modified Genome Assembly

To investigate the woolly locus, two modified OviAri3.1 genome assemblies were produced. To exactly reproduce the locus detailed in [59], the sequence: 'ACTTCTGTAAAATGCAAGATAAATTAAAGTTATTATAACAGTGATTCTTTCAAAAAA ATAAAAACACCATGA' was inserted at position chr25:7,452,231/7,452,232. This modified genome represents the woolly locus. To construct the ancestral locus (i.e. with no asEIFS insertion in chromosome 25), this assembly was edited to replace sequence chr25:7,450,861-7,452,231 with 'TAAAAACACCATGA', as per [59].

All UDG-treated sequencing data was aligned to these modified assemblies in an identical manner as above to produce two bam files. The coverage at the woolly locus was visualised using Integrative Genomics Viewer (IGV) [60], before and after a MAQ filter of 30 was applied. Since there are three copies of EIF2S2 genes annotated on the OviAri3.1 genome assembly (One genuine EIF2S2 gene located on chromosome 13 and two EIF2S2 pseudogenes located on chromosome 7 and chromosome 25), the mapping quality of reads aligned to the insertion are 0 [59].

The ‘breakpoints’ of the woolly locus were defined at positions chr25:7,450,874/5 and chr25:7,452,217/8 of the modified oviAri3.1 assembly. Any reads which straddled either of these sites were deemed evidence of a woolly allele. The ‘breakpoint’ of the ancestral locus was chr25:7,450,861/2. Similarly any read that straddled this point was deemed evidence of the hairy/ancestral allele. Any read attributed to one genotype must only align to one modified assembly and not the other.

**Microscopy analysis**

Images produced by reflected light and transmitted light microscopy reveal a fat layer between the grain and corium, which is characteristic of domestic sheep skin [61,62] (Supplementary Figure 13). The fibres appear mostly unpigmented, with sporadic dark fibres visible (Supplementary Figure 14).

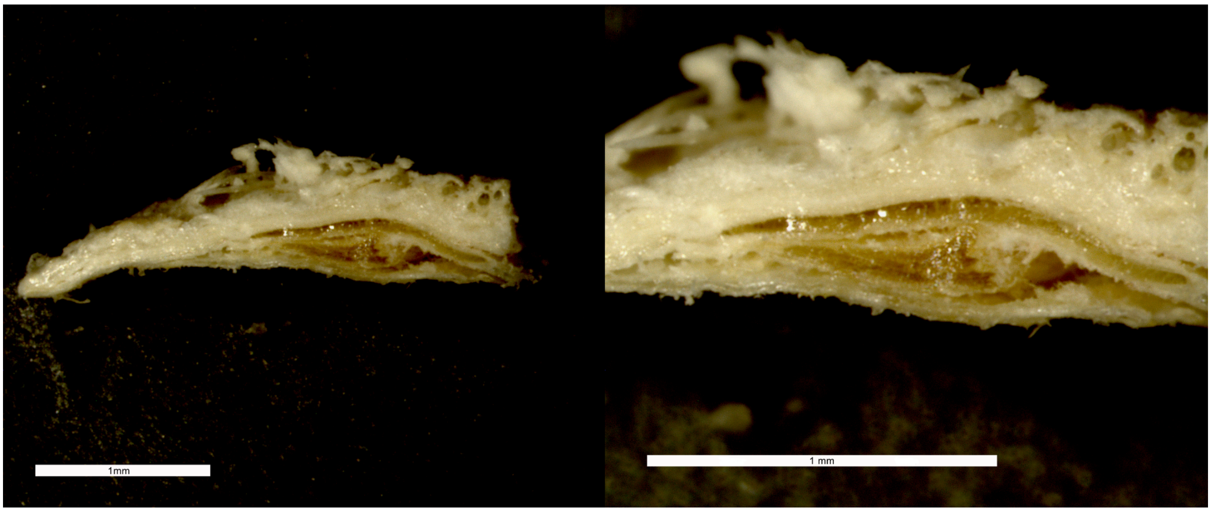


**Supplementary Figure 13: Cross section of the sheep skin. Fat is deposited between the corium layer and the grain.** Images by G. Ruß-Popa.

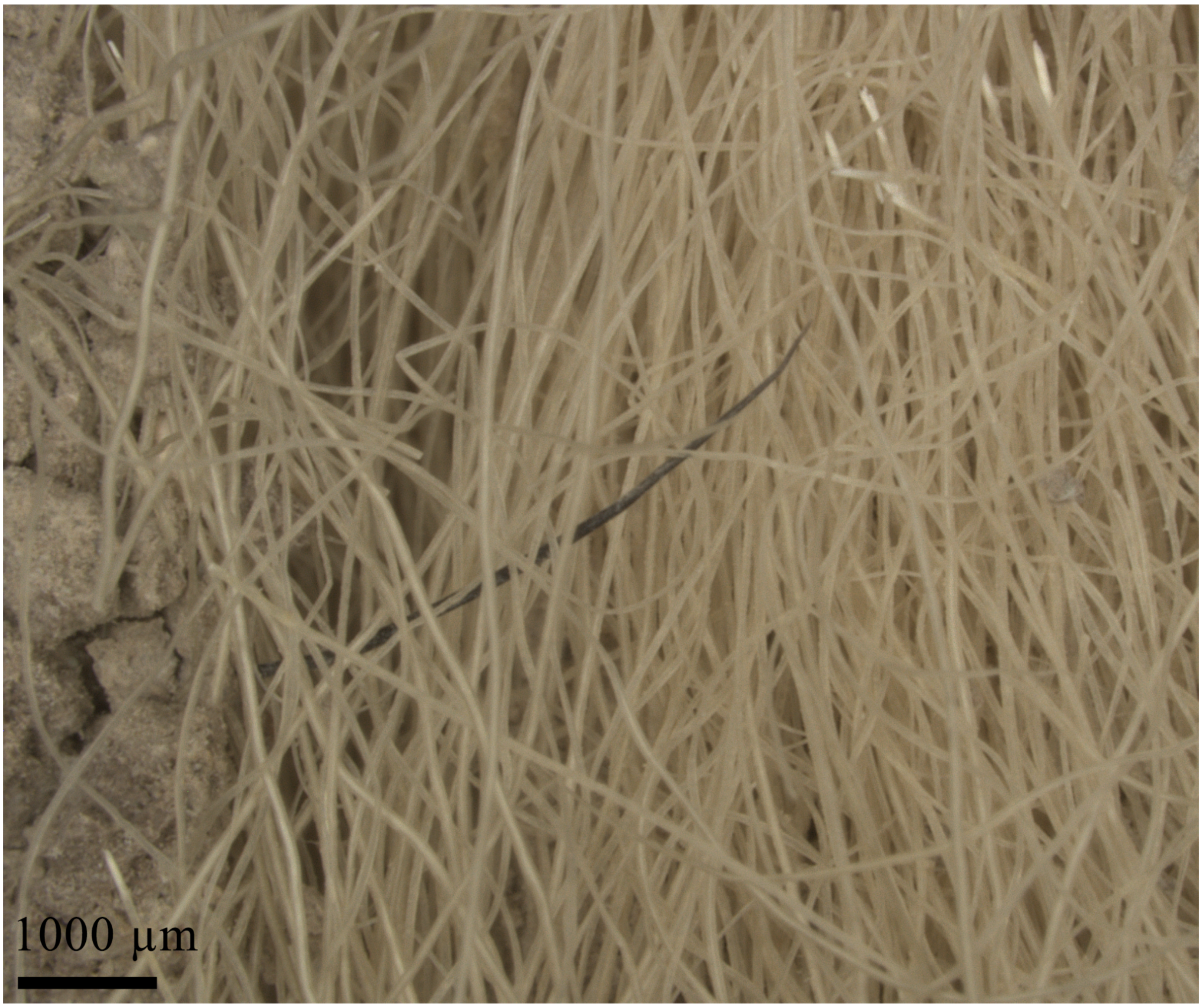


**Supplementary Figure 14**: Microscopy of fibres reveal a mostly unpigmented phenotype. Image by G. Ruß-Popa.

The leg of the sheep is dominated by coarse fibres (primary fibres) (Supplementary Figure 15). Scanning Electron Microscope (SEM) imagery of these hair fibres showed typical structure of sheep hair fibres, with a mosaic type scale and the fine lines on the scale's surface (Supplementary Figure 16A). This is characteristic of sheep hair fibres, and particularly for mouflon and medium-wool breeds. Light microscopy of these coarse fibres revealed an amorphous medulla (Supplementary Figure 16B). Thin unmedullated fibres (underwool) were also detected (Supplementary Figure 17).

Overall, these results are consistent with MUM2 having a hairy/medium-wool phenotype, although results are not conclusive as the fibre types of the lower leg do not reflect the fleece type from the sheep body. Therefore, comparative analysis like wool measurements [63] of different sheep types are planned, in which samples from all parts of the body, including the lower leg, are to be examined.

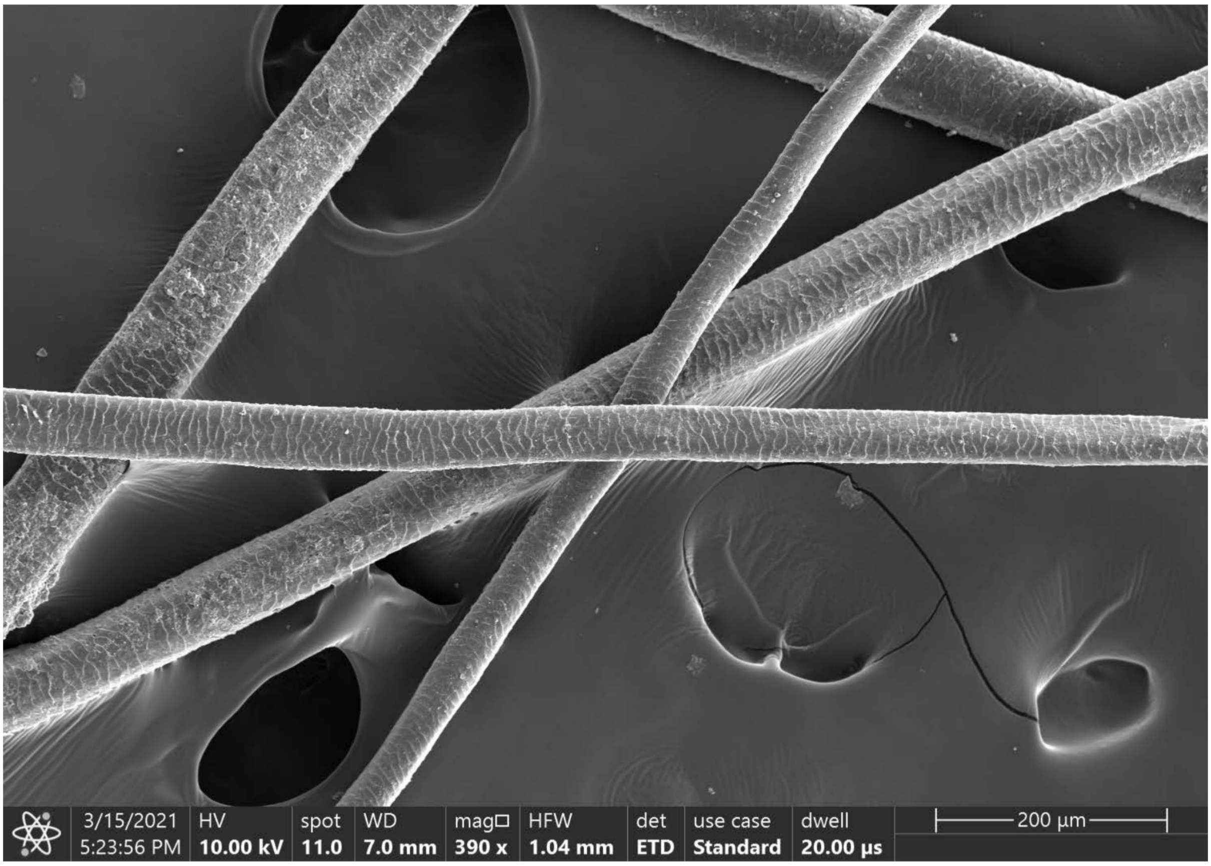


**Supplementary Figure 15: SEM image of different mummy hair fibres.** Image by A. Steiger-Thirsfeld and G. Ruß-Popa.

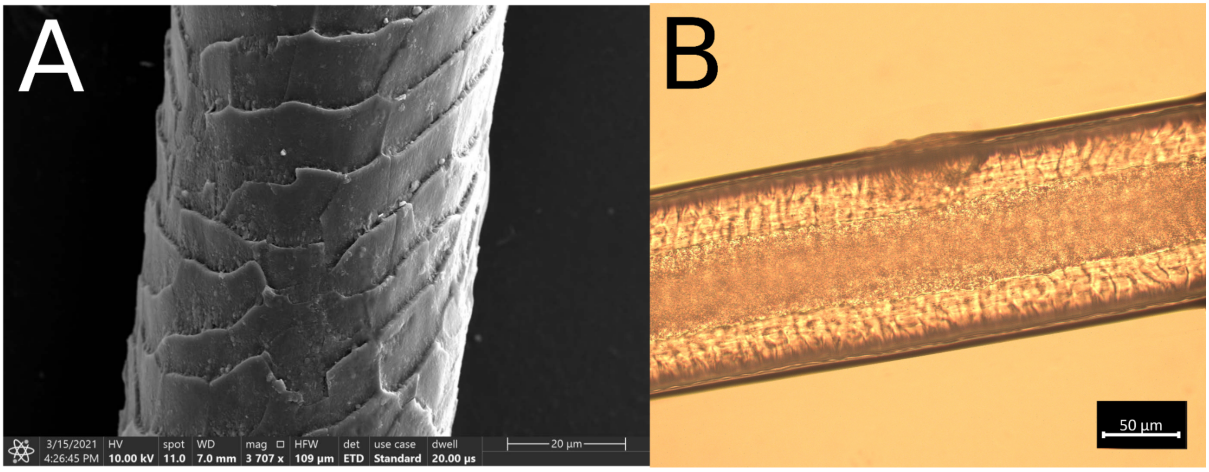


**Supplementary Figure 16A: SEM image of MUM2 hair fibre, displaying the typical mosaic scales of a sheep hair shaft and details of the fine lines on the scale surface.** Image by A. Steiger-Thirsfeld and G. Ruß-Popa. **16B Light microscopy of the coarse hair fibre displaying the amorphous medulla.** Image by G. Ruß-Popa.

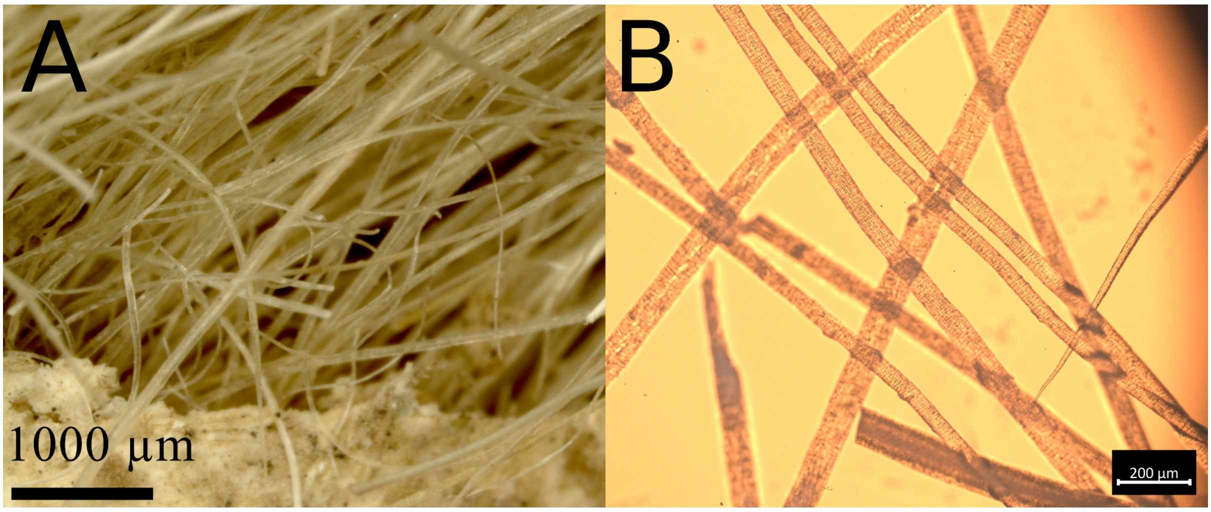


**Supplementary Figure 17A: Microscopy displaying the thin fibres in front.** Image by G. Ruß-Popa. **17B: Light microscopy revealing the unmedullated underwool.** Image by G. Ruß-Popa.

SEM images were generated by G. Ruß-Popa and Andreas Steiger-Thirsfeld at USTEM (Universitäre Service Einrichtung für Transmissions-Elektronenmikroskopie), TU Vienna. FEI Quanta 250 FEGSEM with AFM for Structural investigations (EBSD) was used for SEM imagery. Microscopy was done by G. Ruß-Popa at the Austrian Academy of Sciences, Austrian Archaeological Institute, Archaeological Sciences, Hollandstraße 11-13, 1020 Vienna, Austria. Zeiss SteREO Discovery.V20 (Magnification: 1–150 X) was used for Stereomicroscope (reflected light) analysis. Zeiss Axio Scope.A1 Pol. (Magnification: 1 X – 2.5 X – 5 X – 10 X – 20 X – 40 X) was used for the polarized light microscope analysis for the thin section. Axiocam 305 color was the microscope camera.

(doi:10.1086/basor25066953)

6. Danti MD. 2013 The Late Bronze and Early Iron Age in Northwestern Iran. In *The Oxford Handbook of Ancient Iran* (ed DT Potts), Oxford University Press.

7. Dyson RH, Muscarella OW. 1989 Constructing the Chronology and Historical Implications of Hasanlu IV. *Iran* **27**, 1–27.

8. Danti MD. 2013 Hasanlu V: the Late Bronze and Iron I periods (Hasanlu Excavation Reports 3). *Philadelphia: University of Pennsylvania Museum*

9. Kroll S. 2013 Hasanlu Period III--annotations and corrections. *Iranica Antiqua* **48**, 175–195.

10. Davoudi H. 2017 Subsistence Economy of Iron Age Societies in North-Western Iran Based on Archaeozoological Studies-the Case of Tepe Hasanlu. PhD, Tarbiat Modares University, Iran.

21. Khazaeli R. 2013 The Subsistence Economy of the Old Nishapur from the Formation of the City up to the Mongol Period. MA, University of Tehran.

22. Sambrook J, Fritsch EF, Maniatis T. 1989 *Molecular Cloning: A Laboratory Manual*.

54. Patterson N, Moorjani P, Luo Y, Mallick S, Rohland N, Zhan Y, Genschoreck T, Webster

T, Reich D. 2012 Ancient admixture in human history. *Genetics* **192**, 1065–1093.
